# Genetic and Genomic Analyses of *Drosophila melanogaster* Models of Chromatin Modification Disorders

**DOI:** 10.1101/2023.03.30.534923

**Authors:** Rebecca A. MacPherson, Vijay Shankar, Robert R. H. Anholt, Trudy F. C. Mackay

## Abstract

Switch/Sucrose Non-Fermentable (SWI/SNF)-related intellectual disability disorders (SSRIDDs) and Cornelia de Lange syndrome are rare syndromic neurodevelopmental disorders with overlapping clinical phenotypes. SSRIDDs are associated with the BAF *(Brahma-Related Gene-1* Associated Factor) complex, whereas CdLS is a disorder of chromatin modification associated with the cohesin complex. Here, we used RNA interference in *Drosophila melanogaster* to reduce expression of six genes *(brm, osa, Snr1, SMC1, SMC3, vtd)* orthologous to human genes associated with SSRIDDs and CdLS. These fly models exhibit changes in sleep, activity, startle behavior (a proxy for sensorimotor integration) and brain morphology. Whole genome RNA sequencing identified 9,657 differentially expressed genes (FDR < 0.05), 156 of which are differentially expressed in both sexes in SSRIDD- and CdLS-specific analyses, including *Bap60,* which is orthologous to *SMARCD1,* a SSRIDD-associated BAF component, k-means clustering reveals genes co-regulated within and across SSRIDD and CdLS fly models. RNAi-mediated reduction of expression of six genes co-regulated with focal genes *brm, osa,* and/or Snr1 recapitulated changes in behavior of the focal genes. Based on the assumption that fundamental biological processes are evolutionarily conserved, Drosophila models can be used to understand underlying molecular effects of variants in chromatin-modification pathways and may aid in discovery of drugs that ameliorate deleterious phenotypic effects.

## INTRODUCTION

Switch/Sucrose Non-Fermenting (SWI/SNF)-related intellectual disability disorders (SSRIDDs) and Cornelia de Lange syndrome (CdLS) are syndromic neurodevelopmental Mendelian disorders of chromatin modification. SSRIDDs, including Coffin-Siris syndrome (CSS) and Nicolaides-Baraitser syndrome (NCBRS), stem from variants in genes of the *Brahma-Related Gene-1* Associated Factor (BAF) complex, also known as the mammalian SWI/SNF complex (Hoyer *et al*. 2012; Santen *et al*. 2012; Tsurusaki *et al*. 2012; Van Houdt *et al*. 2012; Tsurusaki *et al*. 2014; Hempel *et al*. 2016; Bramswig *et al*. 2017; Bogershausen and Wollnik 2018; Vasileiou *et al*. 2018; Gazdagh *et al*. 2019; Machol *et al*. 2019; Zawerton *et al*. 2019). CdLS is associated with variants in genes that encode components of the cohesin complex (Krantz *et al*. 2004; Deardorff *et al*. 2007; Deardorff *et al*. 2012; Gil-Rodriguez *et al*. 2015; Boyle *et al*. 2017; Huisman *et al*. 2017; Olley *et al*. 2018). SSRIDD patients exhibit neurodevelopmental delay, intellectual disability, hypotonia, seizures, and sparse hair growth, as well as cardiac, digit, and craniofacial anomalies, where the severity and spectrum of affected phenotypes are dependent upon the specific variant or affected gene product (reviewed in Bogershausen and Wollnik 2018; Schrier Vergano *et al*. 2021; Vasko *et al*. 2021). For example, many SSRIDD patients with variants in *ARID1B* tend to have milder phenotypes including normal growth, milder facial gestalt, and no central nervous system (CNS) abnormalities, whereas most variants in *SMARCB1* are associated with more severe phenotypes, including profoundly delayed developmental milestones, seizures, kidney malformations, and CNS abnormalities (Bogershausen and Wollnik, 2018; Schrier Vergano *et al*. 2021). Furthermore, variants in *ARID1B* are associated with SSRIDD, Autism Spectrum disorder, and non-syndromic intellectual disability (Hoyer *et al*. 2012; De Rubeis *et al*. 2014; Iossifov *et al*. 2014; Vissers *et al*. 2016; van der Sluijs *et al*. 2019). Brain malformations, such as agenesis of the corpus callosum, Dandy-Walker malformation, and cerebellar hypoplasia, have also been observed in 20-30% of all patients with variants in the BAF complex (Vasko *et al*. 2022), but are most commonly observed in patients with variants in *SMARCB1* (Bogershausen and Wollnik 2018).

CdLS patients also display a clinical spectrum including intellectual disability, hirsutism, synophrys, and digit, craniofacial, and CNS anomalies (reviewed in Kline *et al*. 2018; Avagliano *et al*. 2020; Selicorni *et al*. 2021). As in SSRIDDs, some phenotypes are more highly associated with a specific gene, but phenotypic severity can vary widely across variants within the same gene. For example, most patients with variants in *SMC1A* show milder developmental delay and intellectual disability compared to their classical *NIPBL-* CdLS counterparts, but about 40% of *SMC1A* patients exhibit severe epileptic encephalopathy and intellectual disability (Jansen *et al*. 2016; Symonds *et al*. 2017; Selicorni *et al*. 2021).CdLS has also been reclassified as a spectrum of cohesinopathies (Van Allen *et al*. 1993; Kline *et al*. 2018). Patients with pathogenic variants in many genes involved in chromatin accessibility and regulation have overlapping symptoms with CdLS (Parenti et al., 2017; Aoi *et al.* 2019; Cucco *et al.* 2020).

*D. melanogaster* is well-suited for modeling human disorders, as large numbers of flies can be raised economically without ethical or regulatory restrictions. Additionally, SSRIDD- and CdLS-associated genes are highly conserved in flies and a wide variety of genetic tools are available to create fly models of human diseases (Hu *et al*. 2011; Perkins *et al*. 2015; Zirin *et al*. 2020). Previous groups have used *D. melanogaster* to investigate SSRIDDs and CdLS and have observed phenotypes relevant to disease presentation in humans, including changes in sleep, brain function, and brain morphology (Pauli *et al*. 2008; Schuldiner *et al*. 2008; Wu *et al*. 2015; Chubak *et al*. 2019). These studies have provided insight into potential disease pathogenesis and suggested that certain subtypes of SSRIDD and CdLS can be modeled in the fly, but they were not performed in controlled genetic backgrounds.

Here, we present behavioral and transcriptomic data on Drosophila models of SSRIDDs and CdLS in a common genetic background. RNAi-mediated knockdown of Drosophila orthologs of SSRIDD- and CdLS-associated genes show gene- and sex-specific changes in brain structure and sensorimotor integration, as well as increased locomotor activity and decreased night sleep. Transcriptomic analyses show distinct differential gene expression profiles for each focal gene.

## METHODS

### Drosophila Genes and Stocks

We selected SSRIDD-, and CdLS-associated genes with a strong fly ortholog *(Drosophila* RNAi Screening Center Integrative Ortholog Prediction Tool (DIOPT) score > 9) (Hu *et al*. 2011) and a corresponding *attp2* fly line available from the Transgenic RNAi Project (TRiP) (Perkins *et al*. 2015; Zirin *et al*. 2020). We excluded human genes that were orthologous to multiple fly genes to increase the likelihood of aberrant phenotypes upon knockdown of a single fly ortholog. We used *attp40* TRiP lines when assessing phenotypes associated with knockdown of co-regulated genes. We used the *y^1^, sc, v^1^, sev^21^; TRiP2; TRiP3* genotype as the control *UAS* line in all experiments. With the exception of the initial viability screen, we crossed all RNAi lines to a weak ubiquitous *GAL4* driver line, *Ubi156-GAL4* (Garlapow *et al*. 2015). Table S1A lists the Drosophila stocks used.

### Drosophila Culture

For all experiments, we maintained flies at a controlled density on standard cornmeal/molasses medium (Genesee Scientific, El Cajon, CA) supplemented with yeast in controlled environmental conditions (25°C, 50% relative humidity, 12-hour light-dark cycle (lights on at 6 am)). Crosses contained five flies of each sex, with fresh food every 48 hours. After eclosion, we aged flies in mixed-sex vials at a density of 20 flies per vial until used in experiments. We performed experiments on 3-5-day old flies from 8 am to 11 am, unless otherwise noted.

### Viability

For the initial viability screen of Drosophila orthologs of SSRIDD- and CdLS-associated genes, we crossed *attp2* TRiP lines and the control line to three ubiquitous *GAL4* driver lines. For the viability screen of co­regulated genes, we crossed *attp40* TRiP lines and the control line to the *Ubi156-Gal4* driver line. From days 0-15, we noted the developmental stage. For stocks that contained balancers, we recorded the associated phenotypic marker in eclosed progeny.

### Quantitative Real-Time PCR (qRT-PCR)

For the qRT-PCR analyses of gene expression of RNAi targets of *brm, osa, SMC1, SMC3, Snr1,* and *vtd,* we flash froze 3-5-day old whole flies on dry ice and then collected, sexes separately, 30 flies per sample. We stored frozen flies and their extracted RNA at -80°C. We extracted RNA using the Qiagen RNeasy Plus Mini Kit (Qiagen, Hilden, Germany) by homogenizing tissue with 350μL of RLT Plus Buffer containing ²- mercaptoethanol (Qiagen) and DX reagent (Qiagen), using a bead mill at 5m/second for 2 minutes. We quantified RNA with the Qubit RNA BR Assay Kit (ThermoFisher Scientific, Waltham, MA) on a Qubit Fluorometer (ThermoFisher Scientific) according to the manufacturer’s specifications. We synthesized cDNA using iScript Reverse Transcription Supermix (Bio-Rad Laboratories, Inc., Hercules, CA) according to the manufacturer’s instructions. We quantified expression using quantitative real-time PCR with SYBR™ Green PCR Master Mix (ThermoFisher Scientific), according to manufacturer specifications, but with a total reaction volume of 20μL. We used three biological and three technical replicates per sample and calculated percent knockdown using the *ΔΔct* method (Livak and Schmittgen 2001). Table S1B contains primer sequences used. For the qRT-PCR analyses of gene expression for the co-regulated genes *AlplO, CG40485, CG5877, IntS12, Mal-A4,* and *Odd,* we extracted RNA using the Direct-zol RNA MiniPrep Plus Kit (Zymo Research, Irvine, CA) and homogenized tissue with 350μL of Tri-Reagent, using a bead mill at 5m/second for 2 minutes. We used two technical replicates in the qRT-PCR analyses of co­regulated genes.

### Startle-Induced Locomotor Response

We assessed startle response using a variation of a previously described assay (Yamamoto *et al*. 2008). In summary, 36-50 flies per sex per line were placed into individual vials to acclimate 24 hours prior to testing. To standardize the mechanical startle stimulus, we placed a vial housing a single 3-5-day old fly in a chute. Removal of a supporting dowel allows the vial to drop from a height of 4213cm, after which it comes to rest horizontally (Huggett *et al*. 2021). We measured the total time the fly spent moving during a period of 45 s immediately following the drop. We also recorded whether the fly demonstrated a tapping phenotype, a series of leg extensions without forward movement. Time spent tapping was not considered movement for startle calculations.

### Sleep and Activity

We used the Drosophila Activity Monitoring System (DAM System, TriKinetics, Waltham, MA) to assess sleep and activity phenotypes. At 1-2 days of age, we placed flies into DAM tubes containing 2% agar with 5% sucrose, sealed with a rubber cap (TriKinetics) and a small piece of yarn. We collected data for 7 days on a 12-hour light-dark cycle, with sleep defined as at least 5 minutes of inactivity. We discarded data from flies that did not survive the entire testing period, leaving 18-32 flies per sex per line for analysis. We processed the raw sleep and activity data using ShinyR-DAM (Cichewicz and Hirsh 2018) and used the resulting output data for statistical analysis.

### Dissection and Staining of Brains

We dissected brains from cold-anesthetized flies in cold phosphate buffered saline (PBS), before we fixed the brains with 4% paraformaldehyde (v/v in PBS) for 15 minutes, washed with PAXD buffer (1x PBS, 0.24% (v/v) Triton-X 100, 0.24% (m/v) sodium deoxycholate, and 5% (m/v) bovine serum albumin) three times for 10 minutes each, and then washed three times with PBS. We blocked fixed brains with 5% Normal Goat Serum (ThermoFisher Scientific; in PAXD) for 1 hour with gentle agitation, then stained with 2-5 μg/mL of Mouse anti-Drosophila 1D4 anti-Fasciclin II (1:4) (Developmental Studies Hybridoma Bank; Iowa City, IA) for 16-20 hours at 4°C. We washed brains three times with PAXD for 10 minutes and stained them with Goat anti-Mouse IgG-AlexaFluor488 (1:100) (Jackson ImmunoResearch Laboratories, Inc., West Grove, PA) for 4 hours. Then, we washed brains with PAXD three times for 10 minutes each prior to mounting with ProLong Gold (ThermoFisher Scientific). We performed all steps at room temperature with gentle agitation during incubations.

### Brain Measurements

We analyzed 17-20 brains per sex per line using a Leica TCS SPE confocal microscope. We visualized Z-stacks of each brain using Icy v. 2.2.0.0 (de Chaumont *et al*. 2012). We measured ellipsoid body height and ellipsoid body width by measuring vertical ellipsoid body length from dorsal to ventral, and horizontal ellipsoid body length from left to right (relative to the fly). We also measured lengths of the mushroom body alpha and beta lobes by drawing a single 3D line (3DPolyLine Tool within Icy) through the center of each lobe, adjusting the position of the line while progressing through the z-stack. We measured alpha lobes from the dorsal end of the alpha lobe to the alpha/beta lobe heel (where the alpha and beta lobes overlap) and beta lobes from the median end of the beta lobe to the alpha/beta lobe heel. We normalized the measurements for each brain using the distance between the left and right heels of the mushroom body (heel-heel distance). We used the average alpha and beta lobe lengths for each brain for subsequent analyses. In the case of one missing alpha or beta lobe, we did not calculate an average and instead, used the length of the remaining lobe for analysis. If both alpha or both beta lobes were missing, we removed that brain for analysis of the missing lobes, but retained it for analysis of the other brain regions. *We* also recorded gross morphological abnormalities of the mushroom body alpha and beta lobes, including missing lobe, skinny lobe, extra projections, abnormal alpha lobe outgrowth, and beta lobes crossing the midline for each brain. We selected these phenotypes based on prior studies on gross mushroom body morphology (Zwarts *et al.* 2015; Chubak *et al.* 2019).

### Statistical Analyses

Unless noted below, we analyzed all behavioral data and brain morphology data in SAS v3.8 (SAS Institute, Cary, NC) using the “PROC GLM” command according to the Type III fixed effects factorial ANOVA model *Y = μ + L + S + LxS + ε*, where *Y* is the phenotype, *μ* is the true mean, *L* is the effect of line *(e.g.* RNAi line versus the control), *S* is the effect of sex (males, females), and ε is residual error. We performed comparisons between an RNAi line and its control. We also performed additional analyses for each sex separately.

We used a Fisher’s Exact test *(fisher.test* in R v3.63) to analyze the proportion of flies tapping during startle experiments, the number of brains with a specific morphological abnormality, and the number of brains with any gross morphological abnormality.

We performed Levene’s and Brown-Forsythe’s Tests for unequal variances on the same data set used for the analysis of lobe lengths. For both tests, we used the *leveneTest* command *((car* v3.0-ll, Fox and Sanford 2019) in R v3.6.3) to run a global analysis comparing all genotypes as well as pairwise comparisons.

### RNA Sequencing

We synthesized libraries from lOOng of total RNA using the Universal RNA-seq with Nuquant + UDI kit (Tecan Genomics, Inc., CA) according to manufacturer recommendations. We converted RNA into cDNA using the integrated DNase treatment and used the Covaris ME220 Focused-ultrasonicator (Covaris, Woburn, MA) to generate 350bp fragments. We performed ribosomal RNA depletion and bead selection using Drosophila AnyDeplete probes and RNAClean XP beads (Beckman Coulter, Brea, CA), respectively. We purified libraries after 17 cycles of PCR amplification. We measured library fragment sizes on the Agilent Tapestation using the Agilent High Sensitivity DNA 1000 kit (Agilent Technologies) and quantified library concentration using the Qubit 1X dsDNA High Sensitivity Assay kit (Thermo Fisher Scientific). We pooled libraries at 4nM and loaded them onto an lllumina SI flow cell (Illumina, Inc., San Diego, CA) for paired-end sequencing on a NovaSeq6000 (Illumina, Inc., San Diego, CA). We sequenced three biological replicates of pools of 30 flies each per sex per line. We sequenced each sample to a depth of ∼30 million total reads; we resequenced samples with low read depth (<8 million uniquely mapped reads).

We used the default Illumina BaseSpace NovaSeq sequencing pipeline to demultiplex the barcoded sequencing reads. We then merged S1 flow cell lanes, as well as reads from different runs. We filtered out short and low-quality reads using the *AfterQC* pipeline (v0.9.7) (Chen *et al*. 2017) and quantified remaining levels of rRNA via the bbduk command (Bushnell 2014). We aligned reads to the reference genome (D. *melanogaster* v6.13) using GMAP-GSNAP (Wu *et al*. 2016) and counted these unique alignments to Drosophila genes using the featurecounts pipeline from the Subread package (Liao *et al*. 2013). We excluded genes with a median expression across all samples of less than 3 and genes where greater than 25% of the samples had a counts value of 0. We then normalized the data based on gene length and library size using GeTMM (Smid *et al.* 2018) prior to differential expression analysis.

### Differential Expression Analyses

We performed multiple analyses for differential expression in SAS (v3.8; Cary, NC) using the “PROC glm” command. We first performed a fixed effects factorial ANOVA model *Y = μ + L + S + LxS + ε*, where Line *(L,* all RNAi and control genotypes) and Sex (*S*) are cross-classified main effects and LinexSex (LxS) is the interaction term, *Y* is gene expression, *μ* is the overall mean, and ε is residual error. We then performed the same analyses only for genes associated with SSRIDDs or for CdLS; i.e., 9,657 genes that were significantly differentially expressed (FDR < 0.05 for the Line and/or LinexSex terms) in the full model. We ran the ANOVA model for each RNAi genotype compared to the control. Finally, we ran ANOVAs (*Y = μ + L + ε*) separately for males and females for the disease-specific and individual RNAi analyses.

### Gene Ontology and k-means Clustering Analyses

We performed Gene Ontology (GO) statistical overrepresentation analyses on the top 1,000 differentially expressed genes for the Line term (GO Ontology database released 2022-03-22, Pantherdb v16.0 (Mi *et al*. 2013; Thomas *et al*. 2022)) in each disease-specific and pairwise analysis for GO Biological Process, Molecular Function, and Reactome Pathway terms. For the analyses performed on sexes separately, we used the top 600 differentially expressed genes based on the significance of the Line term. The numbers of differentially expressed genes used in GO enrichment gave maximal GO enrichment with minimal redundancy compared to other numbers of differentially expressed genes.

We performed k-means clustering (average linkage algorithm), sexes separately, on Ge-TMM normalized least squares means of 533 genes that had the highest Log2 fold change (FC) in expression. We identified the cutoff threshold value for Log2FC by first sorting genes in a descending order of maximal absolute value of Log2FC, then fitted lines to roughly linear segments of the generated distribution and designated the cutoff threshold as the Log2FC value of the index at the intersection of the two fitted lines. We used hierarchical clustering (Average Linkage algorithm, WPGMA) to determine the approximate number of natural clusters, then performed clustering with varying values of k to determine the largest number of unique, but not redundant, expression patterns. We also performed GO statistical overrepresentation analyses on genes in each k-means cluster (GO Ontology database released 2022-07-01, Pantherdb v17.0 (Mi *et al*. 2013; Thomas *et al*. 2022)) in each disease-specific and pairwise analysis for GO Biological Process, Molecular Function, and Reactome Pathway terms.

## RESULTS

### Drosophila Models of SSRIDDs and CdLS

We identified Drosophila orthologs of 12 human genes associated with the SSRIDD chromatin remodeling disorders and CdLS with a DIOPT score > 9 and for which TRiP RNAi lines in a common genetic background and without predicted off-target effects were publicly available. Using these criteria, the Drosophila genes *Baplll, brm, osa,* and *Snr1* are models of SSRIDD-associated genes *ARID1A, ARID1B, SMARCA2, SMARCA4, SMARCB1,* and *SMARCE1;* and *Nipped-B, SMC1, SMC3,* and *vtd* are models of CdLS-associated genes *NIPBL, SMC1A, SMC3,* and *RAD21* (Table S2). We obtained L/AS-RNAi lines generated in the same genetic background for each of the fly orthologs and crossed these RNAi lines to each of three ubiquitous *GAL4* drivers to assess viability (Figure S1). We selected ubiquitous drivers since the human SSRIDD- and CdLS-associated genes and Drosophila orthologs are ubiquitously expressed, and SSRIDD and CdLS patients carry pathogenic variants in all cells. We initially crossed each L/AS-RNAi line to three ubiquitous *GAL4* drivers *(Actin-GAL4, Ubiquitin-GAL4,* and *Ubi156-GAL4)* and assessed viability and degree of gene knockdown in the F1 progeny (Figure SI). *Ubiquitin-GAL4-med’\ated* gene knockdown resulted in viable progeny in only three of the eleven L/AS-RNAi lines, with most progeny dying during the embryonic or larval stage (Figure S1). Based on these data, we selected the weak ubiquitin driver *Ubi156-GAL4* (Garlapow *et al*. 2015) and the *UAS-RNAi* lines for *brm, osa, Snr1, SMC1, SMC3,* and *vtd* for further study (Table 1). With the exception of *Ubi156>osa* males which had ∼15% gene knockdown, RNAi knockdown of all genes ranged from 40-80% (Table S3). Given that SSRIDDs and CdLS are largely autosomal dominant disorders, knockdown models that retain some degree of gene expression are reflective of the genetic landscape of SSRIDD and CdLS patients.

**Table 1.** Drosophila genes used in fly models. The table indicates fly genes used in SSRIDD and CdLS fly models, as well as the respective human orthologs and MIM numbers, associated human disease and respective MIM numbers, and DIOPT scores. Human orthologs are only included in the table if the DIOPT score is greater than 9.

| Fly Gene | Human Ortholog(s) | Human Ortholog MIM number(s) | Associated Human Disease | Phenotype MIM Number(s) | DIOPT score |
| --- | --- | --- | --- | --- | --- |
| *brm* | *SMARCA2, SMARCA4* | 600014, 603254 | SSRIDD (NCBRS, CSS 4) | 601358, 614609 | 13, 12 |
| *osa* | *ARID1A, ARID1B* | 603024, 614556 | SSRIDD (CSS 2, CSS 1) | 614607, 135900 | 12, 12 |
| *SMC1* | *SMC1A* | 300040 | Cornelia de Lange syndrome 2 | 300590 | 12 |
| *SMC3* | *SMC3* | 606062 | Cornelia de Lange syndrome 3 | 610759 | 12 |
| *Snr1* | *SMARCB1* | 601607 | SSRIDD (CSS 3) | 614608 | 15 |
| *vtd* | *RAD21, RAD21L1* | 606462, 619533 | Cornelia de Lange syndrome 4 | 614701 | 11, 10 |

**Table 2.** Differentially expressed gene counts. The table shows the number of differentially expressed genes (FDR < 0.05) for the Line and/or Line x Sex terms for each pairwise analysis of knockdown vs control, sexes together and sexes separately.

| Comparison | Analysis |  |  |  |
| --- | --- | --- | --- | --- |
|  | Both sexes |  | Females only | Males only |
|  | Line | Line×Sex | Line | Line |
| *brm* vs. Control | 2808 | 1652 | 583 | 2995 |
| *osa* vs. Control | 2179 | 1059 | 1135 | 1580 |
| *Snr1* vs. Control | 4996 | 3632 | 3026 | 3376 |
| *SMC1* vs. Control | 2714 | 1727 | 2540 | 2395 |
| *SMC3* vs. Control | 1874 | 586 | 2711 | 1161 |
| *vtd* vs. Control | 1998 | 961 | 818 | 1630 |

### Effects on Startle Response

Given the neurological and musculoskeletal clinical findings in SSRIDD, and CdLS patients (Bogershausen and Wollnik 2018; Kline *et al*. 2018; Avagliano *et al*. 2020; Schrier Vergano *et al*. 2021; Selicorni *et al*. 2021; Vasko *et al*. 2022), we assessed startle-induced sensorimotor integration for RNAi of *brm, osa, Snr1, SMC1, SMC3,* and *vtd* relative to their control genotype. Almost all genotypes exhibited a decreased startle response across both sexes *(p <* 0.02 for all by-sex by-genotype comparisons to the control, Figure 1A, Table S4). Males with *osa* or *brm* knockdown did not exhibit changes in startle response *(p >* 0.05), and females with *Snr1* knockdown showed an increased startle response *(p <* 0.0001). In the lines where both sexes were affected, we observed more extreme phenotypes in males (Figure 1A).

**Figure 1.**
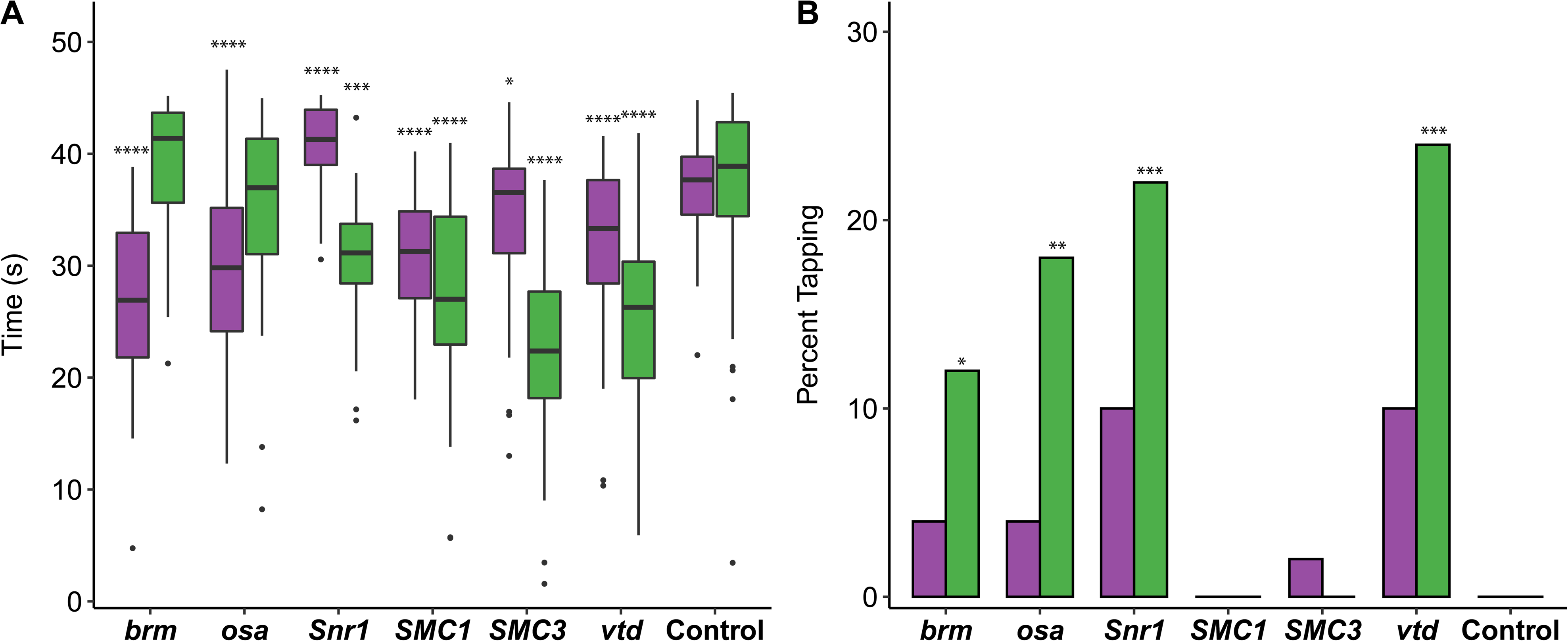
Altered startle response phenotypes in SSRIDD and CdLS fly models. Startle phenotypes of flies with *Ubil56-GAL4-med’\ated* RNAi knockdown. (A) Boxplots showing the time, in seconds, spent moving after an initial startle force. Asterisks represent sex-specific pairwise comparisons with the control. (B) Bar graphs showing the percentage of flies that exhibit tapping behavior (see File SI and S2) following an initial startle stimulus. Females and males are shown in purple and green, respectively. See Table S4 for ANOVAs (A) and Fisher’s Exact Tests (B). N - 36-50 flies per sex per line. *: *p* < 0.05, **: *p <* 0.01, ***: p < 0.001, ****: *p <* 0.0001.

While testing flies for startle response, we noticed that some flies exhibited a specific locomotion phenotype we termed “tapping”. Tapping is characterized by repetitive extension and retraction of individual legs as if to walk, but without progressive movement in any direction (File S1). Compared to the control (example shown in File S2), we observed an increase in the number of flies exhibiting tapping behavior in male flies with knockdown of *brm (p =* 0.0267), *osa (p =* 0.0026), *Snrl (p -* 0.0005) and *vtd (p* = 0.0002) (Figure 1B, Table S4). We also observed increases in tapping behavior in females with knockdown of *Snrl* and *vtd* that fall just outside of a significance level of 0.05 *(p =* 0.0563 for both genes); Figure 1B, Table S4). The tapping and startle phenotypes were not evident across all genes associated with a specific disorder.

### Effects on Sleep and Activity

We hypothesized that hypotonia and sleep disturbances observed in SSRIDD and CdLS patients (Liu and Krantz 2009; Stavinoha *et al*. 2011; Rajan *et al*. 2012; Zambrelli *et al*. 2016; Bogenshausen and Wollnik 2018; Schrier Vergano *et al*. 2021; Vasko *et al*. 2021) may correspond to changes in activity and sleep in Drosophila models. Sleep disturbances were also observed in a previous Drosophila model of *NIPBL-* CdLS (Wu *et al*. 2015). Therefore, we quantified activity and sleep phenotypes for RNAi-mediated knockdown of *brm, osa, Snr1, SMC1, SMC3,* and *vtd.* All RNAi genotypes showed increases in overall spontaneous locomotor activity *(p <* 0.02 for all by-sex by-genotype comparisons to the control, Figure 2A, Table S4). This increase in spontaneous locomotor activity was most pronounced in males with knockdown of *osa (p <* 0.0001); this was the only genotype for which males were more active than females (Figure 2A, Table S4). All RNAi genotypes showed decreases in night sleep *(p <* 0.0001 for all by­sex by-genotype comparisons to the control). Flies with knockdown of *osa* (males, *p <* 0.0001; females, *p* < 0.0001) and females with knockdown of *vtd (p <* 0.0001) spent about half of the nighttime awake, the least amount of sleep across all flies tested (Figure 2B, Table S4). In addition to increased activity, the Drosophila models of SSRIDDs and CdLS have fragmented sleep: the number of sleep bouts at night was increased for all lines and sexes compared to the control *(p <* 0.0001 for all by-sex by-genotype comparisons to the control, except *SMC1* males, *p =* 0.0023, Figure 2C, Table S4).

**Figure 2.**
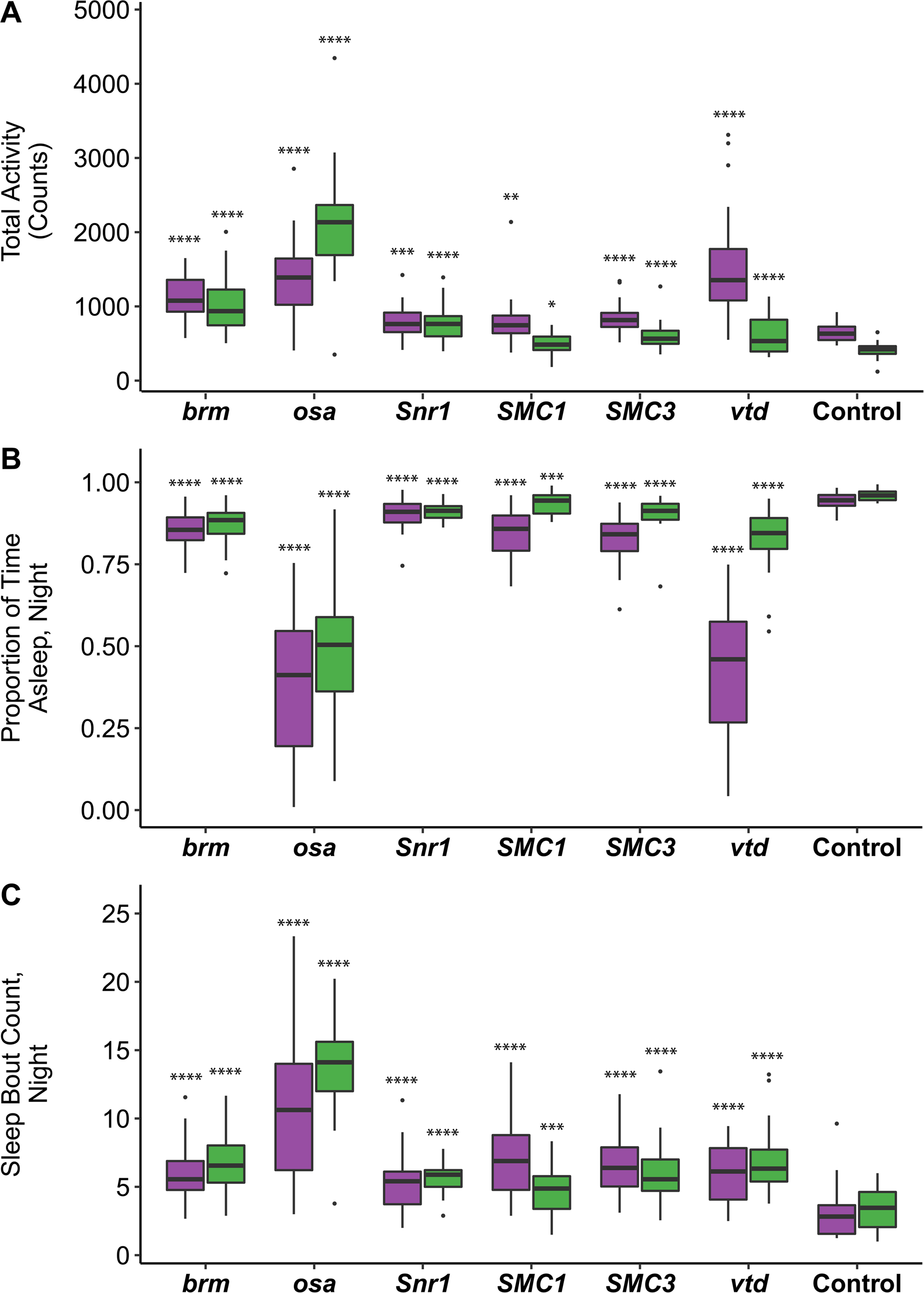
Altered sleep and activity phenotypes in SSRIDD and CdLS fly models. Boxplots displaying activity and sleep phenotypes of flies with *Ubil56-GAL4-med’\ated* RNAi knockdown. (A) total activity; (B) proportion of time spent asleep at night; (C) number of sleep bouts at night. Females and males are shown in purple and green, respectively. N - 18-32 flies per sex per line. See Table S4 for ANOVAs. Asterisks indicate pairwise comparisons of each line to the control, sexes separately. *: *p* < 0.05, **: *p <* 0.01, ***: *p* < 0.001, ****: *p <* 0.0001.

### Effects on Brain Morphology

To assess changes in brain structure in *brm, osa, Snr1, SMC1, SMC3,* and *vtd* RNAi genotypes, we focused on the mushroom body and the ellipsoid body, as prior studies on SSRIDDs in flies have shown changes in mushroom body structure (Chubak *et al*. 2019), and the mushroom body has been linked with regulation of sleep and activity in Drosophila (Joiner *et al.* 2006; Pitman *et al*. 2006; Guo *et al.* 2011; Sitaraman *et al.* 2015). Furthermore, SSRIDD and CdLS patients often present with intellectual disability and CNS abnormalities (Bogershausen and Wollnik 2018; Kline *et al*. 2018; Avagliano *et al*. 2020; Schrier Vergano *et al*. 2021; Selicorni *et al*. 2021; Vasko *et al*. 2022). In the Drosophila brain, the mushroom body mediates experience-dependent modulation of behavior (reviewed in Modi *et al*. 2020), making the mushroom body and the ellipsoid body, which mediates sensory integration with locomotor activity, suitable targets for examining changes in brain structure. We used confocal microscopy to quantify the lengths of both alpha and beta lobes of the mushroom body, as well as the horizontal and vertical lengths of the ellipsoid body (Figures 3A-B). The lengths of these lobes were measured in three dimensions, capturing the natural curvature of the alpha and beta lobes of the mushroom body instead of relying upon a 2D measurement of a 3D object.

**Figure 3.**
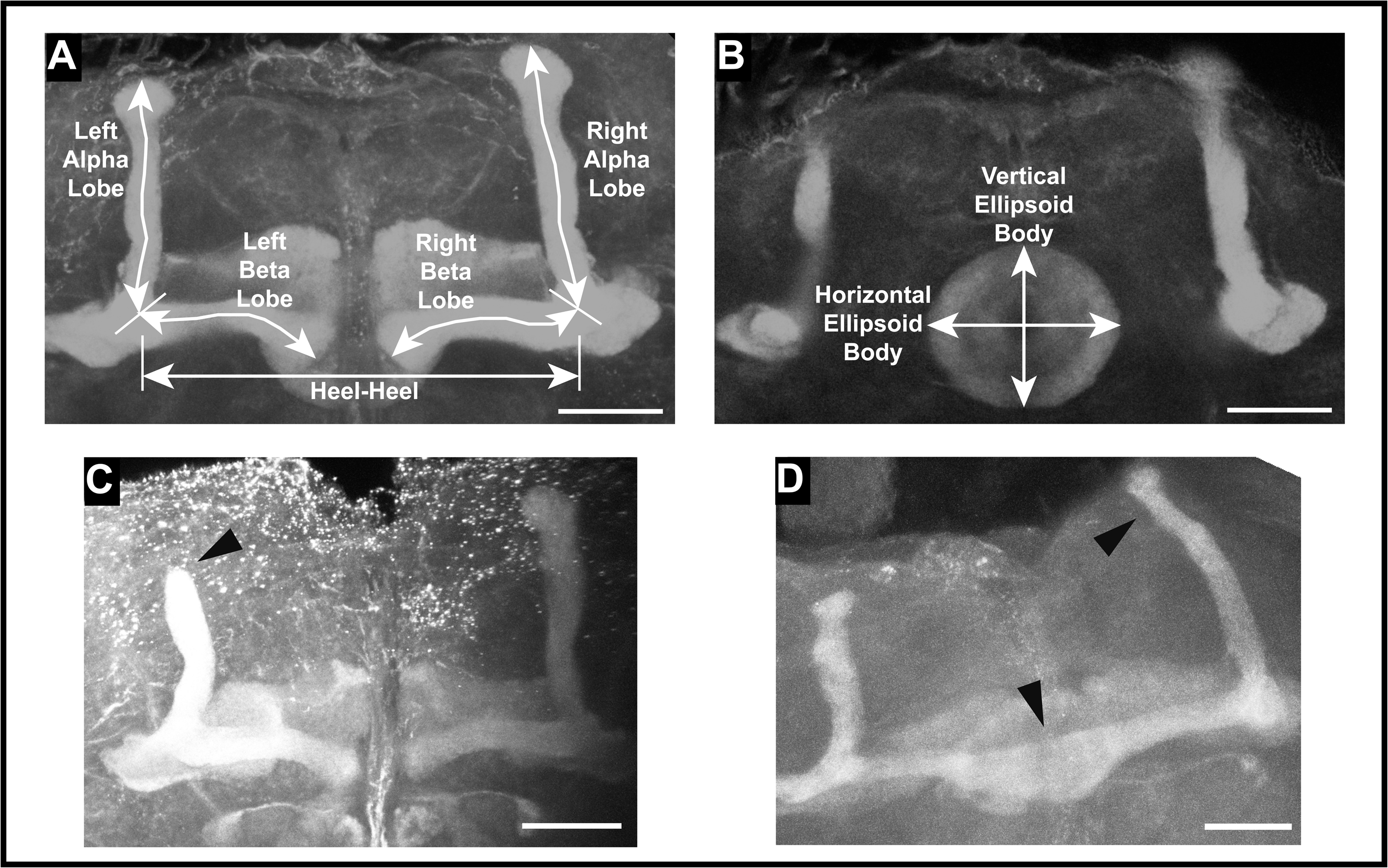
Examples of mushroom body abnormalities in SSRIDD and CdLS fly models. Images of a wild type mushroom body annotated with measurement descriptors for (A) mushroom body alpha and beta lobes, and heel-heel normalization measurement; and (B) ellipsoid body measurements. Images of select brains from flies with *Ubil56-GAL4-med’\ated* RNAi knockdown of *osa* showing (C) stunted alpha lobe outgrowth and narrowed alpha lobe head in a female oso-deficient fly brain; and (D) beta lobe crossing the midline/fused beta lobes, as well as a skinny alpha lobe in a male oso-deficient fly brain. Images shown are z-stack maximum projections from confocal imaging. Triangular arrowheads indicate the abnormalities. The scale bar represents 25 pM.

We observed sex-specific changes in brain morphology (Figure 3C-D). Females, but not males, showed decreased ellipsoid body dimensions with knockdown of *Snr1* (horizontal, *p =* 0.0002; vertical, *p <* 0.0444, Table S4), while knockdown of *vtd* in females showed decreased alpha *(p =* 0.0088) and beta *(p =* 0.0433) lobe lengths. In addition to sex-specific effects, we observed sexually dimorphic effects; females with knockdown of *brm* showed decreases in alpha lobe and horizontal ellipsoid body length *(p -* 0.0409, *p =* 0.0224, respectively), while *brm* knockdown males showed increases in alpha lobe and horizontal ellipsoid body length *(p =* 0.0301, *p =* 0.0305, respectively; Figure 4, Table S4). Levene’s tests for equality of variances indicate that the ellipsoid body measurements have sex-specific unequal environmental variances in some genotypes compared to the control (Figure 4, Table S4). These results show that these models of SSRIDDs and CdLS show morphological changes in the mushroom body and ellipsoid body.

**Figure 4.**
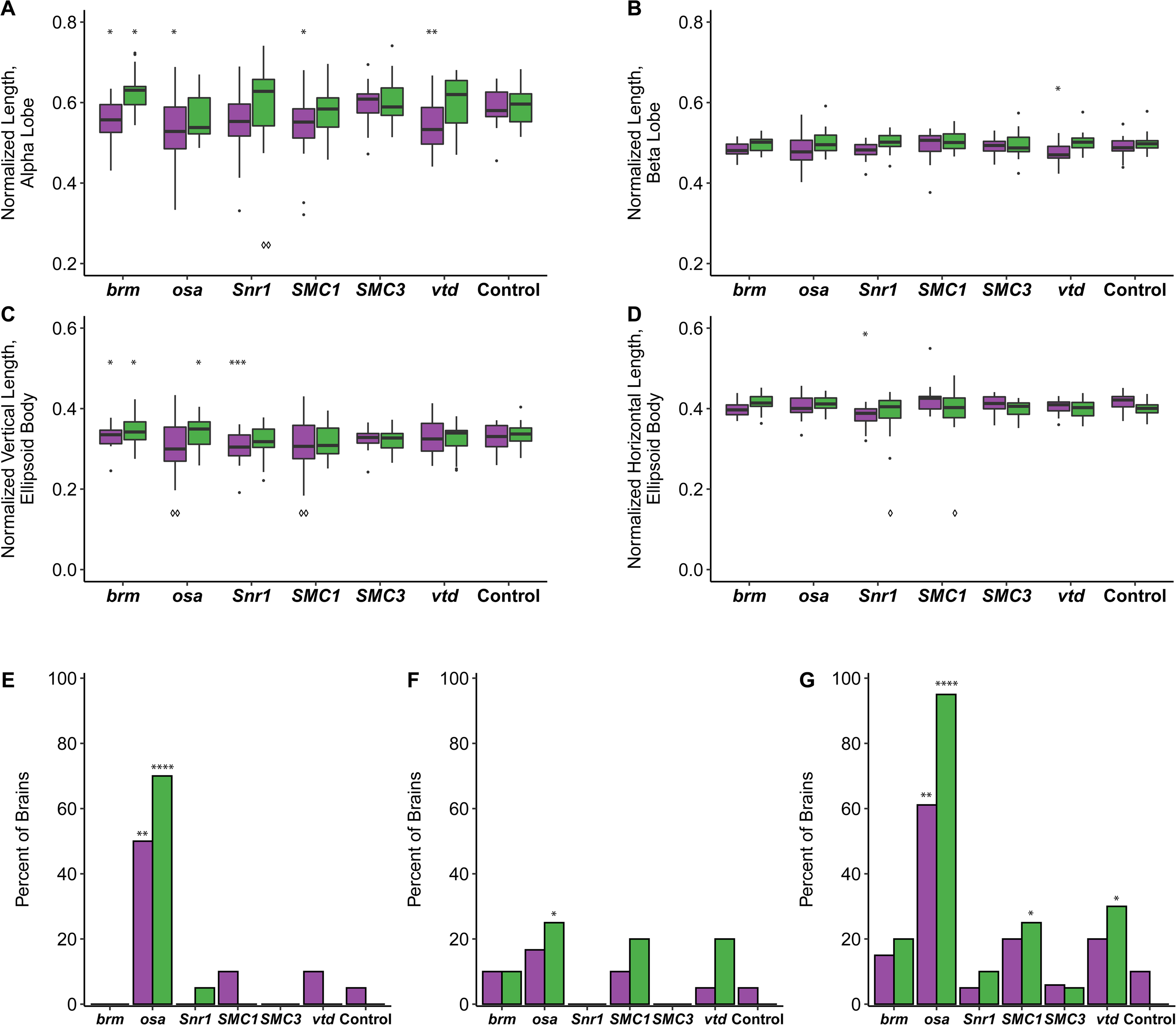
SSRIDD and CdLS fly models show gene-specific changes in mushroom body and ellipsoid body. Boxplots showing (A) the average alpha lobe and (B) beta lobe length for each brain; (C) ellipsoid body height (vertical direction; dorsal-ventral) and (D) width (left-right; lateral). Bargraphs showing the percentage of brains that (E) have a stunted alpha lobe(s)/narrowed alpha lobe head(s); (F) have a beta lobe(s) crossing the midline, including fused beta lobes; and (G) display one of more of the following defects: skinny alpha lobe, missing alpha lobe, skinny beta lobe, missing beta lobe, stunted alpha lobe/narrowed alpha lobe head, beta lobe crossing the midline/fused beta lobes, extra projections off of the alpha lobe, extra projections off of the beta lobe. See Figure 3. All brains were dissected from flies with *Ubil56-GAL4-med’\ated* RNAi knockdown. For panels A-D, brains missing only one alpha or beta lobe are represented by the length of the remaining lobe and brains missing both alpha lobes or both beta lobes were not included in the analyses. For panels E-G, data were analyzed with a Fisher’s Exact test, sexes separately. Asterisks (*) and diamonds (panels A-D only; 0) represent pairwise comparisons of the knockdown line versus the control in ANOVAs or Fisher’s Exact tests, and Levene’s tests for unequal variances, respectively. See Table S4 for ANOVAs, Fisher’s Exact and Levene’s Test results. Females and males are shown in purple and green, respectively. N - 17-20 brains per sex per line. *: *p <* 0.05, **: *p* < 0.01, ***: *p <* 0.001, ****: *p <* 0.0001. 0: *p* < 0.05, 00: *p <* 0.01.

We also recorded gross morphological abnormalities, such as missing lobes, beta lobes crossing the midline, and impaired/abnormal alpha lobe outgrowth (Figure 3C-D). Although each abnormality was observed across multiple genotypes, only flies with knockdown of *osa* demonstrated consistent brain abnormalities. Male and female *osa* knockdown flies both exhibited an increased number of alpha lobes with impaired outgrowth (males: *p <* 0.0001, females: *p <* 0.0025, Figure 4E, Table S4), and the *osa* knockdown males also showed a significant number of beta lobe midline defects *(p =* 0.0471, Figure 4F, Table S4). Males with knockdown of *SMC1* and *vtd* also showed increased numbers of abnormal brains *(p -* 0.0471, *p =* 0.0202 respectively; Figure 4G, Table S4). Changes in brain morphology are more gene-and sex-dependent than changes in sleep, activity, and startle response.

### Effects on Genome-wide Gene Expression

We performed genome-wide analysis of gene expression for the *brm, osa, Snr1, SMC1, SMC3,* and *vtd* RNAi genotypes and their control, separately for males and females. We first performed a factorial fixed effects analysis of variance (ANOVA) for each expressed transcript, partitioning variance in gene expression between sexes, lines, and the line by sex interaction for all seven genotypes. We found that 8,481 and 6,490 genes were differentially expressed (FDR < 0.05 for the Line and/or LinexSex terms, Table S5), for a total of 9,657 unique genes.

*brm, osa, Snr1* and their human orthologs (Tables 1, S2) are part of the same protein complex (BAF complex in humans, BAP-complex in flies). Therefore, we evaluated whether other BAP complex members *Bap55, Bap60,* and *Baplll* (which are orthologous to human BAF complex members *ACTL6A, SMARCD1,* and *SMARCE1,* respectively), are differentially expressed in the analysis of all genes. We observed differential expression of strong fly orthologs (DIOPT > 9) of additional BAF complex subunits in the global model and found that *Bap55* and *Bap60* (FDR-corrected Line p-values: 0.0123, 0.01306, respectively; Table S5), but not *Baplll,* are differentially expressed. We did not observe differential expression of *Nipped-B* in the global analysis. *Nipped-B* is a member of the fly cohesin complex along with *SMC1, SMC3,* and *vtd,* and is orthologous to the human cohesin complex member *NIPBL*.

We next performed separate pairwise analyses for SSRIDD-associated fly orthologs and CdLS-associated fly orthologs against the control genotype using the subset of 9,657 unique differentially expressed genes from the full ANOVA model (Tables 2, S5). We also performed these analyses on sexes separately (Tables 2, S5). The number of differentially expressed genes at a given FDR threshold varies across pairwise comparisons and across sexes. For example, females with knockdown of *brm* and *Snr1* have 583 and 3,026 differentially expressed genes (FDR < 0.05), respectively, whereas males with knockdown of these genes have 2,996 and 3,376 differentially expressed genes (FDR < 0.05), respectively (Tables 2, S5). We observed the largest number of differentially expressed genes in flies with knockdown of Snr1 (Tables 2, S5). At FDR < 0.0005, there were still 1,059 genes differentially expressed in *Snr1* males (Table S5). A greater number of differentially expressed genes are upregulated than downregulated in flies with knockdown of *brm, SMC1, SMC3,* and *vtd* (Table S5). In contrast, flies with knockdown of *osa* and *Snr1* have a greater number of downregulated genes (Table S5). Flies with knockdown of Snr1 and *SMC1* had the greatest percentage of differentially expressed genes shared between males and females: 12.2% (698) and 7.6% (348) respectively (Table S6). *Snr1* also had the greatest percent knockdown by RNAi. Only four genes are differentially expressed in all pairwise comparisons of knockdown lines versus the control line, in both males and females; all are computationally predicted genes (Table S6). *We* performed k-means clustering to examine patterns of co-regulated expression, separately for males (k=S) and females (k=10). We identified the cutoff threshold value for Log2FC by first sorting genes in a descending order of maximal absolute value of Log2FC (Table S7). We fitted lines to roughly linear segments of the generated distribution and designated the cutoff threshold as the Log2FC value of the index at the intersection of the two fitted lines (Figure S2, Table S7). The genes in each cluster are listed in Table S8. Although many clusters reveal gene-specific expression patterns *(e.g.* Cluster F1, F9, F10, Figure 5; Clusters M1, M6, Figure 6), Clusters F7 and F8 show disease-specific patterns, where knockdown of form, *osa,* and *Snr1* clusters separately from *SMC1, SMC3,* and *vtd* (Figure 5). This is not surprising, as form, *osa,* and *Snr1* are part of the fly BAF complex and models for SSRIDDs, whereas *SMC1, SMC3,* and *vtd* are associated with the fly cohesin complex and are models for CdLS. We also observed patterns involving genes from both SSRIDDs and CdLS. Clusters F4 and M3 contain genes upregulated in response to knockdown of *SMC3, osa,* and form and downregulated in response to knockdown of Snr1 and *SMC1* (Figures 5-6) Clusters F5 and M5 contain genes upregulated only in flies with knockdown of *osa* and *Snr1* (Figures 5-6). Notably, many long noncoding RNAs (lncRNAs) feature prominently in many of the male and female clusters (Figures 5-6; Tables S7, S8).

**Figure 5.**
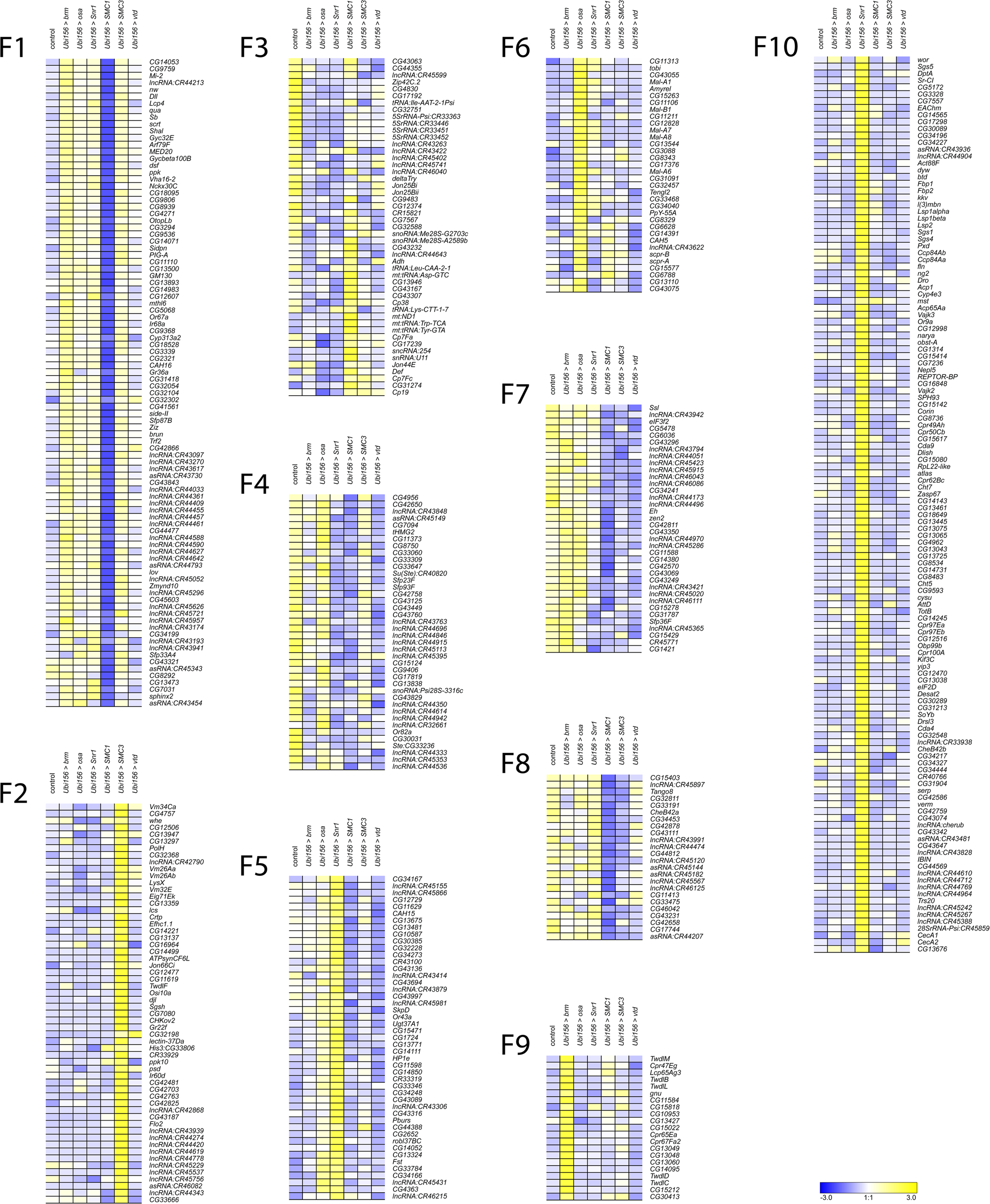
k-means clusters for females. k-means clusters (k = 10, average linkage algorithm) based on expression patterns of the 535 genes with maximal absolute value of the fold-change in expression, compared to the control. Blue and yellow indicate lower and higher expression, respectively.

**Figure 6.**
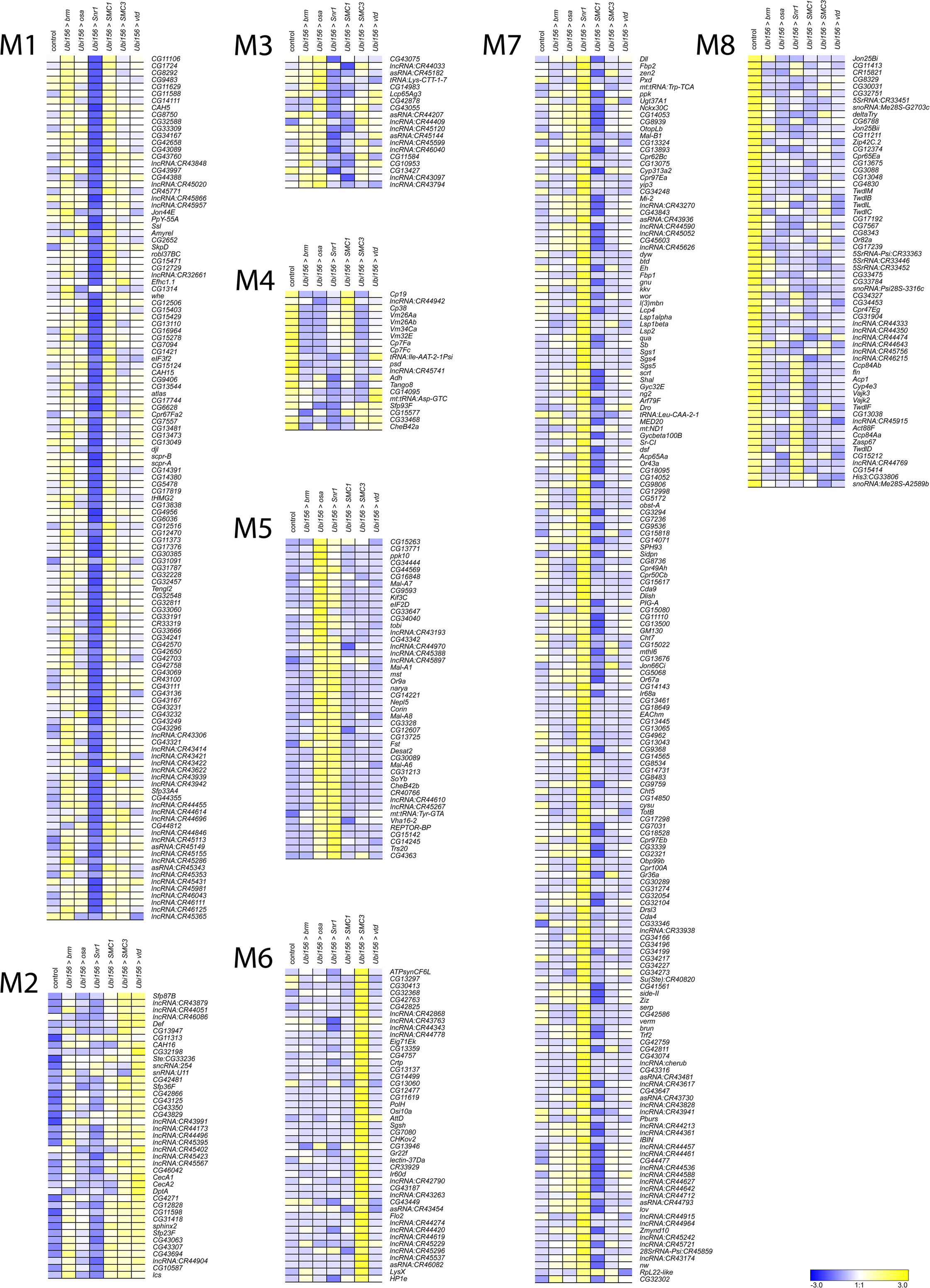
k-means clusters for males. k-means clusters (k *= 8,* average linkage algorithm) based on expression patterns of the 535 genes with maximal absolute value of the fold-change in expression, compared to the control. Blue and yellow indicate lower and higher expression, respectively.

To infer functions of these differentially expressed genes, we performed Gene Ontology (GO) analyses on the top approximately 600 (1000) differentially expressed genes for sexes separately (sexes pooled) (Table S9). These analyses reveal that differentially expressed genes associated with knockdown of CdLS-associated fly orthologs are involved in chromatin organization, regulation and processing of RNA, reproduction and mating behavior, peptidyl amino acid modification, and oxidoreductase activity (Table S9). We also see sex-specific effects, such as muscle cell development in males and neural projection development in females (Table S9). Differentially expressed genes associated with knockdown of SSRIDD-associated fly orthologs in males are involved in mating behavior, cilia development, and muscle contraction, while we see overrepresented ontology terms involved in chromatin modification, mitotic cell cycle, and serine hydrolase activity in females (Table S9). We observed more alignment of GO terms across genes and sexes in the CdLS fly models *(SMC1, SMC3, vtd)* than in SSRIDD fly models (form, *osa, Snrl).* There were no overrepresented GO terms for females in the CdLS-specific analysis. However, in the 156 genes shared across both sexes and both the SSRIDD and CdLS disease-level analyses, we see an overrepresentation of muscle cell development and actin assembly and organization (Table S9). GO enrichment on k-means clusters does not reveal over-representation of any biological processes, molecular functions or pathways for Clusters F7, F8, F4, F5, and M3 (Table S10). Genes involved in alpha-glucosidase activity are overrepresented in Cluster M5 (Table S10).

We generated Venn diagrams (Figure S3) to display the degree of similarity in differentially expressed genes across analyses, including the 156 genes shared across SSRIDD and CdLS males and females (Table S6). Interestingly, 93% (2689/2907) of genes differentially expressed in a disease-specific analysis of CdLS males were also differentially expressed in CdLS females or in SSRIDD fly models (Table S6). This is in contrast to CdLS females, SSRIDD males, and SSRIDD females, in which about 25% of the differentially expressed genes were specific to a single analysis (Table S6). Approximately 24 and 56 percent of the differentially expressed genes (FDR<0.05) in pairwise comparisons for males and females, respectively, have a predicted human ortholog (DIOPT > 9) (Table Sil).

### Co-Regulated Genes

We selected a subset of co-regulated genes from gene expression analyses as potential modifiers of the focal genes form, *osa,* and/or *Snr1. We* chose genes that had a significant effect (Line FDR < 0.05) in analyses pooled across sexes, a suggestive effect (Line FDR < 0.1) for each sex separately, a greater than or less than two-fold-change in both sexes, a strong human ortholog (DIOPT > 9), and an available offp40TRiP RNAi line (the same genetic background as the focal genes). We increased the FDR threshold to 0.1 for the sex-specific pairwise analyses to account for the decreased power of these analyses compared to those with sexes combined. This resulted in 31 genes (Table S12). We further narrowed our selection by prioritizing genes for further study with potential roles in neurological tissues, metabolism, chromatin, orthologs associated with disease in humans, and computationally predicted genes of unknown function. The six fly genes we selected for further study are *AlplO, CG40485, CG5877, IntS12, Mal-A4,* and *Odd,* which are orthologous to human genes *ALPG, DHRS11, NRDE2, INTS12, SLC3A1,* and *ODC1,* respectively (human ortholog with highest DIOPT score listed; Table S12). All six genes tested were co-regulated with *Snr1,* but *CG40485* and *CG5877* were not co-regulated with *osa* and *brm* models of SSRIDDs (Table S6).

For each target gene, we crossed the *UAS-RNAi* line to the *Ubi156-GAL4* driver and performed qRT-PCR to assess the magnitude of reduction in gene expression. All co-regulated genes had reduced expression in both sexes (Table S13). We then assessed the effects of these genes on startle response, sleep, and activity. Knockdown of *Mal-A4, CG5877* and *AlplO* showed changes in startle response times for both sexes (Figure S4A, Table S14). *Mal-A4* demonstrated sexually dimorphic changes in startle response similar to flies with *Snr1* knockdown, as females showed an increase *(p =* 0.0215) and males showed a decrease *(p <* 0.0001) in startle response (Figure S4A, Table S14). We also quantified tapping behavior in these co-regulated genes and found that flies with knockdown of *CG5877* and *Odd* showed an increase in tapping behavior compared to the control, similar to flies with knockdown of *osa* and *Snrl* (Figure IB), although we only observed tapping in females with knockdown of *Odd* (Figure S4B, Table S14; *CG5877* females: *p =* 0.0266, *CG5877* males: *p <* 0.0001; *Odd* females: *p =* 0.0125). With the exception of *CG40485,* which showed no changes in sleep or activity for either sex, all male RNAi genotypes had increased nighttime sleep bouts *(p <* 0.03), decreased night sleep *(p <* 0.03), and, with the additional exception of *CG5877* RNAi flies, increased overall activity *(p <* 0.006) (Figure S4, Table S14). Knockdown of *Mal-A4* and *Odd* also showed increased activity for females *(p =* 0.0049, *p -* 0.0044, respectively). Only knockdown of *CG5877* resulted in increased night sleep for females *(p =* 0.0014) (Figure S4C-D, Table S14). These changes in activity and sleep phenotypes largely parallel those observed for SSRIDD fly models (Figure 2, Table S14).

Based on effects on startle response, tapping behavior, locomotor activity, night sleep, and sleep bouts, none of the phenotypes associated with RNAi of the co-regulated genes exactly matched the phenotypes associated with RNAi of the SSRIDD focal genes in both magnitude and direction. However, three genes *(Mal-A4, CG5877, Odc1)* exhibited at least one altered phenotype in both sexes (Figure S4). These phenotypic observations provide evidence that *Mal-A4, CG5877,* and/or *Odc1* may be interacting with the focal genes of the SSRIDD fly models.

## DISCUSSION

Variants in members of the mammalian SWI/SNF complex (BAF complex) give rise to SSRIDDs, Mendelian disorders with a wide range of phenotypic manifestations, including Coffin-Siris and Nicolaides-Baraitser syndromes (reviewed in Bogershausen and Wollnik 2018; Schrier Vergano *et al*. 2021). The diverse consequences of such variants and variation in penetrance of similar variants in different affected individuals suggest the presence of segregating genetic modifiers. Such modifiers may represent targets for ameliorating therapies or serve as indicators of disease severity, yet they cannot be easily identified in humans due to the limited sample size of individuals with rare disorders. In addition to identifying potential modifiers, Drosophila models can be used to understand underlying molecular effects of variants in chromatin-modification pathways and may aid in discovery of drugs that ameliorate deleterious phenotypic effects.

We used a systematic comparative genomics approach to generate Drosophila models of disorders of chromatin modification, based on the assumption that fundamental elements of chromatin modification are evolutionarily conserved. First, we reduced expression of BAF and cohesin complex orthologs through targeted RNA interference with a *GAL4* driver that induces minimal lethality. We assessed consequences of target gene knockdown on behaviors that mimic those affected in patients with SSRIDDs and CdLS. We used startle behavior, a proxy for sensorimotor integration, and sleep and activity phenotypes to assess the effects of variants in fly orthologues of human genes associated with similar behavioral disorders. These Drosophila models show increased activity, decreased night sleep, and changes in sensorimotor integration. Although we cannot readily recapitulate cognitive developmental defects in Drosophila, these behavioral phenotypes along with brain morphology measurements provide a representative spectrum of behaviors that correlate with human disease phenotypes. We observed gene-specific effects. In addition to showing the largest changes in sleep and activity phenotypes, only *osa* RNAi flies showed stunted mushroom body alpha lobes. Furthermore, only females with knockdown of *Snr1* showed an increase in startle response times. Our neuroanatomical studies focused on morphological changes in the ellipsoid body and mushroom bodies. We cannot exclude effects on other regions in the brain.

Next, we performed whole genome transcriptional profiling to identify co-regulated genes with each focal gene and used stringent filters to identify candidate modifier genes from the larger subset of co­regulated genes, k-means clustering reveals co-regulated genes unique to knockdown of a single protein complex member (Figures S4, S5), yet also shows genes co-regulated in response to knockdown of several, but not all, members of the fly cohesin and SWI/SNF complexes. Gene-specific and cross-disease effects are intriguing, since *brm, osa,* and *Snr1* are part of the fly SWI/SNF complex, and *SMC1, SMC3,* and *vtd* are part of the fly cohesin complex, yet have widespread gene-specific downstream effects on gene regulation. Upon knockdown of one protein complex member, we did not necessarily find changes in gene expression of other members of the same complex. It is possible that a compensatory mechanism exists that maintains transcript levels of other fly SWI/SNF or cohesin complex members or the focal genes themselves (Dorsett 2009; Raab *et al*. 2017; Van der Vaart *et al*. 2020), such as with *Nipped-B* in a CdLS fly model (Wu *et al*. 2015). Furthermore, the abundance of lncRNAs co-regulated with focal genes (Figures S4, S5, Table S8) is intriguing given the association between lncRNAs, chromatin modification, and changes in gene expression in both flies and humans (Li *et al.* 2019; Statello *et al.* 2021). *Snr1* is part of the Brahma complex, a core component of the BAP complex and is orthologous to *SMARCB1* (Table S2). *Odd,* which encodes ornithine decarboxylase, is orthologous to *ODC1* (Table S12), which is associated with Bachmann-Bupp syndrome, a rare neurodevelopmental disorder with alopecia, developmental delay, and brain abnormalities (Prokop *et al*. 2021; Bupp *et al*. 2022). Ornithine decarboxylase is the rate-limiting step of polyamine synthesis, which provides critical substrates for cell proliferation and differentiation (reviewed in Wallace *et al.* 2003; Pegg 2016). Polyamines interact with nucleic acids and transcription factors to modulate gene expression (Watanabe *et al*. 1991; Hobbs and Gilmour 2000; Miller-Fleming *et al*. 2015; Maki *et al*. 2017). *CG5877* is predicted to mediate post­transcriptional gene silencing as part of the spliceosome (Herold *et al*. 2009) and is orthologous to human *NRDE2* (Table S12). *Mal-A4* is predicted to be involved in carbohydrate metabolism (Inomata *et al*. 2019) and is orthologous to *SLC3A1* (Table S12). We observed extensive sexual dimorphism in behavioral phenotypes and transcriptional profiles upon knockdown of SSRIDD- and CdLS-associated genes. Although we are not aware of transcriptional profiles currently available for SSRIDD patients, RNA sequencing of post-mortem neurons from CdLS patients have shown dysregulation of hundreds of neuronal genes (Weiss *et al*. 2021). RNA sequencing in a A//’pped-6-mutation fly model of *NIPBL-CdiS* found differential expression of “2800 genes in the imaginal disc (FDR < 0.05) (Wu *et al*. 2015). Thus, we believe the number of differentially expressed genes upon gene knockdown reported herein is comparable to previous studies.

## DATA AVAILABILITY

All high throughput sequencing data are deposited in GEO GSE213763. Raw behavioral data, qPCR data, and coding scripts are available on GitHub at https://github.com/rebeccamacpherson/Dmelmodels CSS NCBRS CdLS. All L/AS-RNAi lines used in this study are available at the Bloomington Drosophila Stock Center, except the ubiquitous RNAi driver *Ubi156-GAL4* and the double RNAi lines, which are available upon request.

## ACKNOWLEDGEMENTS

We thank Dr. Lakshmi Sunkara for assistance with RNA sequencing, Marion R. Campbell III, Miller Barksdale, and Rachel C. Hannah for technical assistance with behavioral assays and brain dissections. We thank Dr. Joshua Walters for helping create Figure 3 and helping dissect brains, and Dr. Richard Steet at the Greenwood Genetic Center for suggestions. We thank Katelynne Collins and Tori Gyorey for assistance with the RNAi studies. We thank the TRiP at Harvard Medical School (NIH/NIGMS R01- GM084947) for providing transgenic RNAi fly stocks used in this study.

## AUTHOR CONTRIBUTIONS

RAM performed all experiments. VS assisted with RNA sequencing analysis; TFCM conceptualized the research program and TFCM and RRHA directed the research program. TFCM, RRHA, and RAM provided resources and wrote the manuscript.

## SUPPLEMENTARY INFORMATION

Figure S1. **Gross viability observations in potential CSS/NCBRS and CdLS fly models.** Life stage shown is the final stage of the *Drosophila* life cycle where live individuals were observed. “X” indicates flies did not have detectable levels of gene knockdown, as quantified via qRT-PCR.

Figure S2. **Selection of genes for k-means clustering.** Elbow plots of maximal fold change in expression plotted against rank order (blue) across all analyses for each of 9657 genes (A) and for genes with a maximum fold change difference greater than 4 (B). See Table S7. The red and green lines were fit to roughly linear segments of the generated distribution (blue). The orange lines are drawn from the plot elbow (determined by the x coordinate of the intersection of the green and red lines) to the x and y axes.

Figure S3. **Overlap of differentially expressed genes in SSRIDD and CdLS fly models.** Venn diagrams displaying the number of differentially expressed genes (FDR < 0.05), in **SSRIDD** and CdLS fly models, sexes separately. Pairwise gene-specific analyses from (A) CdLS fly models and (B) **SSRIDD** fly models. Panel (C) shows overlap of disease-specific analyses, pooled across disease-associated genes.

Figure S4. **Altered phenotypes due to knockdown of co-regulated genes.** Bar plots displaying differences in the average values of the experimental line versus the control line for (A) startle response, (B) percent of flies tapping, (C) total activity, and (D) proportion of time asleep at night. All lines have *Ubi156-GAL4-med’\ated* RNAi knockdown. Females and males are shown in purple and green, respectively. See Table S14 for ANOVAs (A,B,D) and Fishers Exact Tests (C). N=29-32 per sex per line. Error bars represent standard error of the difference based on error propagation (Burns and Dobson 1981). Asterisks represent pairwise analyses of the experimental line vs the control, sexes separately. *: *p <* 0.05, **: *p <* 0.01, ***: *p <* 0.001.

File S1. **Video of tapping behavior in a male fly with knockdown of *vtd* following a startle response.**

File S2. **Video of control male fly following a startle response.**

Table S1. **Fly reagents and primer sequences.** Drosophila reagents and primer sequences. (A) Drosophila lines used. (B) Primer sequences used for qRT-PCR. BDSC: Bloomington Drosophila Stock Center.

Table S2. **Ortholog prediction scores for potential focal genes.** Human-Drosophila ortholog prediction scores generated using Drosophila RNAi Screening Center Integrative Ortholog Prediction Tool (DIOPT). Human genes associated with SSRIDDs and Cornelia de Lange syndrome.

Table S3. **Percent knockdown of focal genes.** Average RNAi-mediated qRT-PCR knockdown of focal genes.

Table S4. **Quantification of changes in behavior and brain morphology from knockdown of focal genes.** Quantification of changes in behavior and brain morphology from RNAi knockdown. Statistical analyses characterizing SSRIDD and CdLS fly models. (A) ANOVAs for startle response. (B) Fisher’s Exact Tests for tapping behavior. (C) ANOVAs for sleep and activity measurements. (D) ANOVAS for mushroom body lobe lengths. (E) Levene’s and Brown-Forsythe Tests for unequal variances of mushroom body lobe length data. (F) Gross brain abnormalities. Line and Sex are fixed effects, df: degrees of freedom, SS: Type III Sum of Squares, MS: Mean Squares.

Table S5. **ANOVA results from differential expression analyses.** Gene name, gene symbol, FlyBase ID, normalized read counts (counts per million), and raw and Benjamini-Hochberg FDR adjusted p-values for all genes for all model terms used in the ANOVA analyses. (A) Full model using all knockdown lines and the control according to the model *Y = μ + Line + Sex + Line x Sex +* ? for 15915 genes. (B-G) Pairwise comparisons of single gene knockdown vs. the control (sexes together *Y = μ + Line + Sex + Line x Sex +* ?; and sexes separately *Y = μ + Line +* ?) on the 9657 genes from the full model differentially expressed (FDR < 0.05) for the *Line* and/or *Line x Sex* terms. (B) *brm.* (C) *osa.* (D) *Snr1.* (E) *SMC1.* (F) *SMC3.* (G) *vtd.* (H-I) Disease-specific comparisons (sexes together *Y = μ + Line + Sex + Line x Sex +* ?; and sexes separately *Y = μ. + Line +* ?). (H) SSRIDDs. (I) Cornelia de Lange syndrome (CdLS).

Table S6. **Overlap of differentially expressed genes across analyses.** FDR-corrected p-values less than 0.05 for the Line term of each of the 9657 genes. (A) Pairwise analyses of each knockdown line compared to the control, sexes separately. (B) Disease-specific analyses, sexes separately. NA indicates FDR-corrected P-values for the effect of Line greater than 0.05.

Table S7. **k-means threshold.** (A) Average log2 fold change values for each differentially expressed gene for each set of samples, as well as maximum, minimum across all samples. (B) Determination of threshold by ranking, indexing and fitting lines to fold change plots, fc: Iog2 fold change; f: females, m: males.

Table S8. **k-means clustering gene lists.** Lists of genes within each k-means cluster. (A) Females. (B) Males.

Table S9. **Gene Ontology (GO) analyses for differentially expressed genes.** “Analysis” indicates the gene set used in the analysis.

Table S10. **Gene Ontology (GO) analyses for k-means clusters.** “Analysis” indicates the gene set used in the analysis.

Table S11. **Ortholog prediction scores for differentially expressed genes**. Drosophila-human ortholog prediction scores, generated using Drosophila RNAi Screening Center Integrative Ortholog Prediction Tool (DIOPT). Differentially expressed fly genes for each by-sex pairwise comparison.

Table S12. **Ortholog prediction scores and known disease associations for co-regulated genes.** Drosophila-human ortholog prediction scores, generated using Drosophila RNAi Screening Center Integrative Ortholog Prediction Tool (DIOPT) and Online Mendelian Inheritance of Man (OMIM)-derived known disease/phenotype associations and corresponding MIM numbers. Subset of 31 Drosophila genes co-regulated with brm, osa, and/or Snrl.

Table S13. **Percent knockdown of co-regulated genes.** Average RNAi-mediated qRT-PCR knockdown of co-regulated genes.

Table S14. **Quantification of changes in behavior from knockdown of co-regulated genes.** Quantification of changes in behavior from RNAi knockdown of co-regulated genes. (A) ANOVAs for startle response. (B) Fisher’s Exact Tests for tapping behavior. (C) ANOVAs for sleep and activity measurements. Line and Sex are fixed effects, df: degrees of freedom, SS: Type III Sum of Squares, MS: Mean Squares.

