## Supplementary figures and images for "Genetic and Genomic Analyses of *Drosophila melanogaster* Models of Chromatin Modification Disorders"

### Supplemental Figure 1

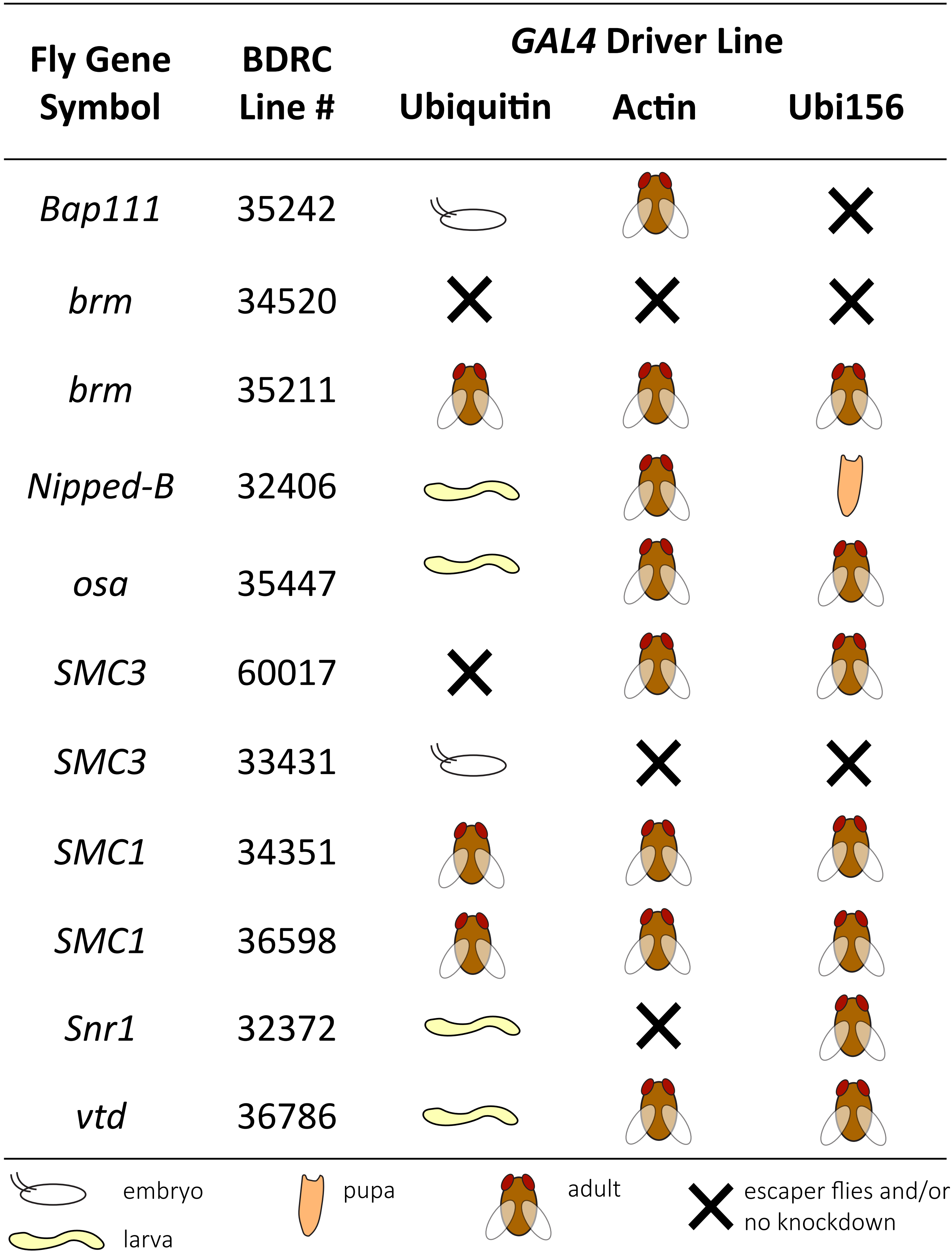

### Supplemental Figure 2

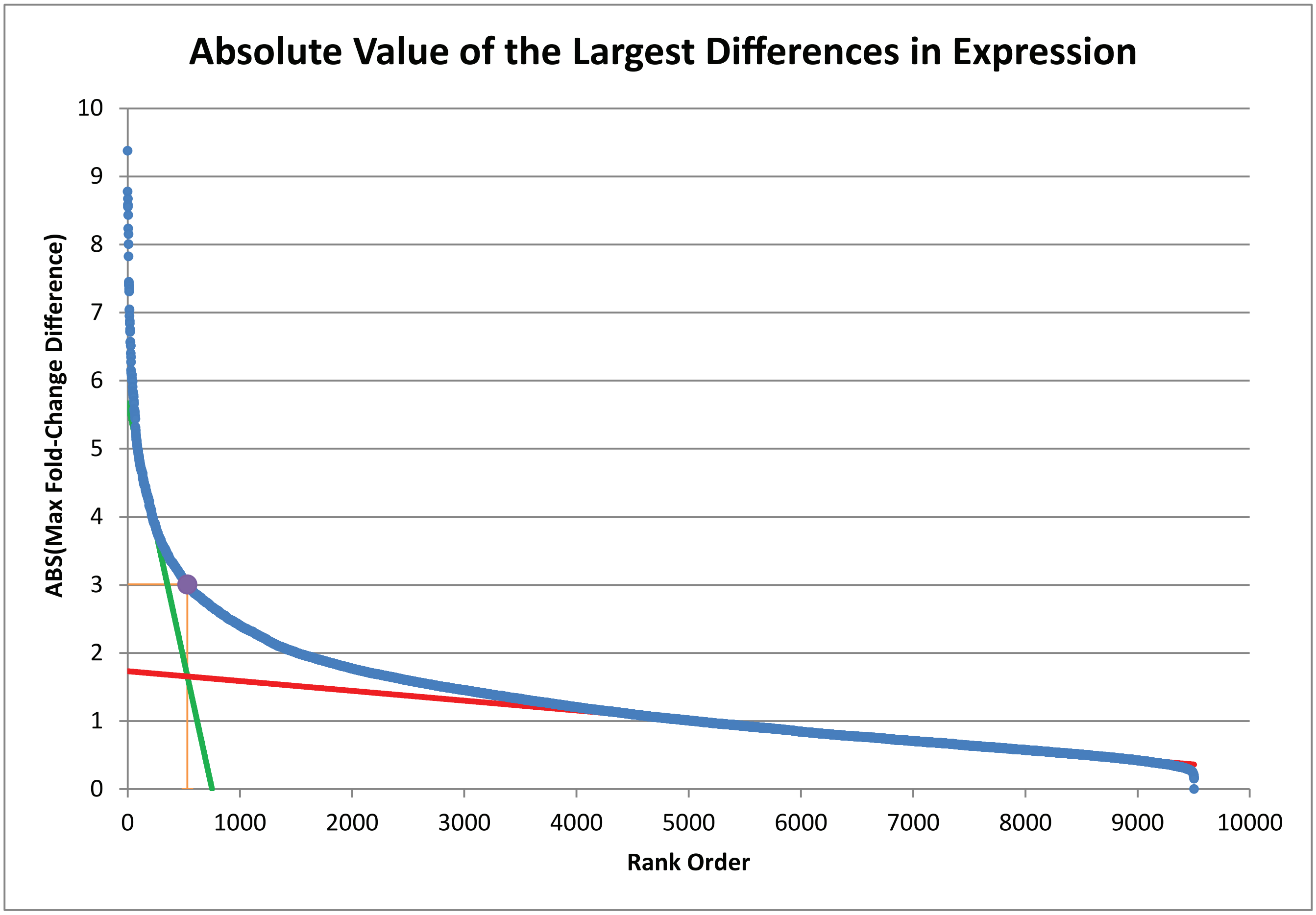

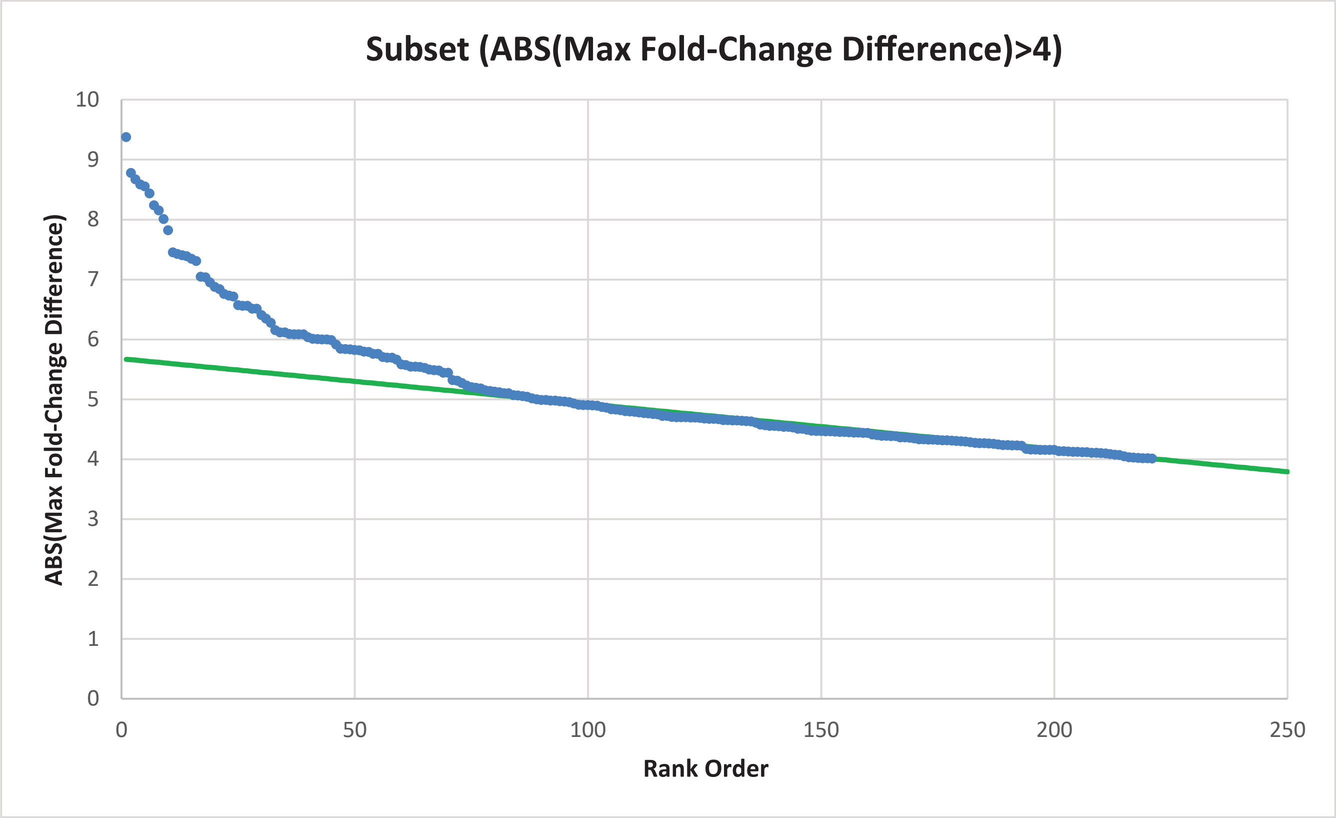

### Supplemental Figure 3

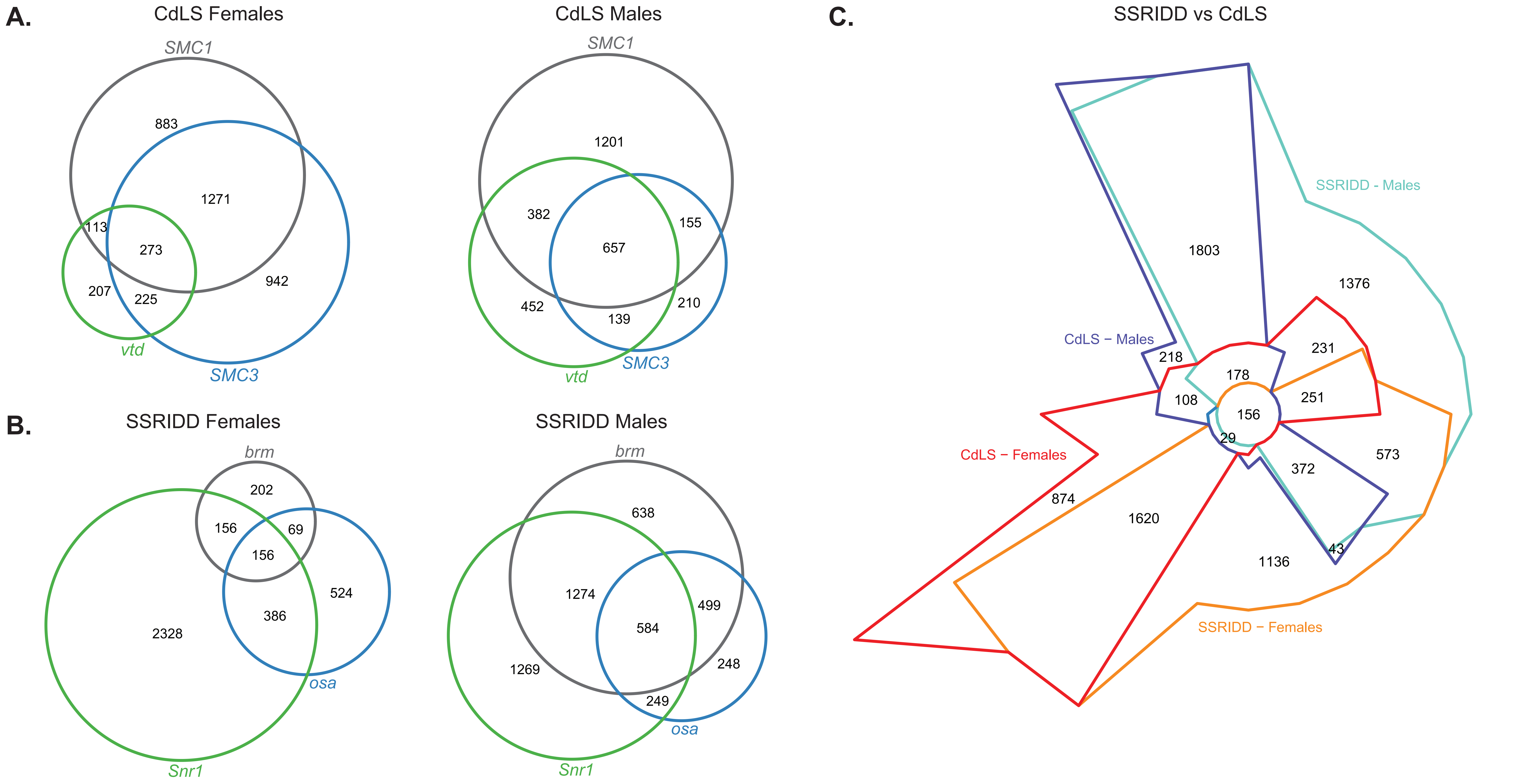

### Supplemental Figure 4

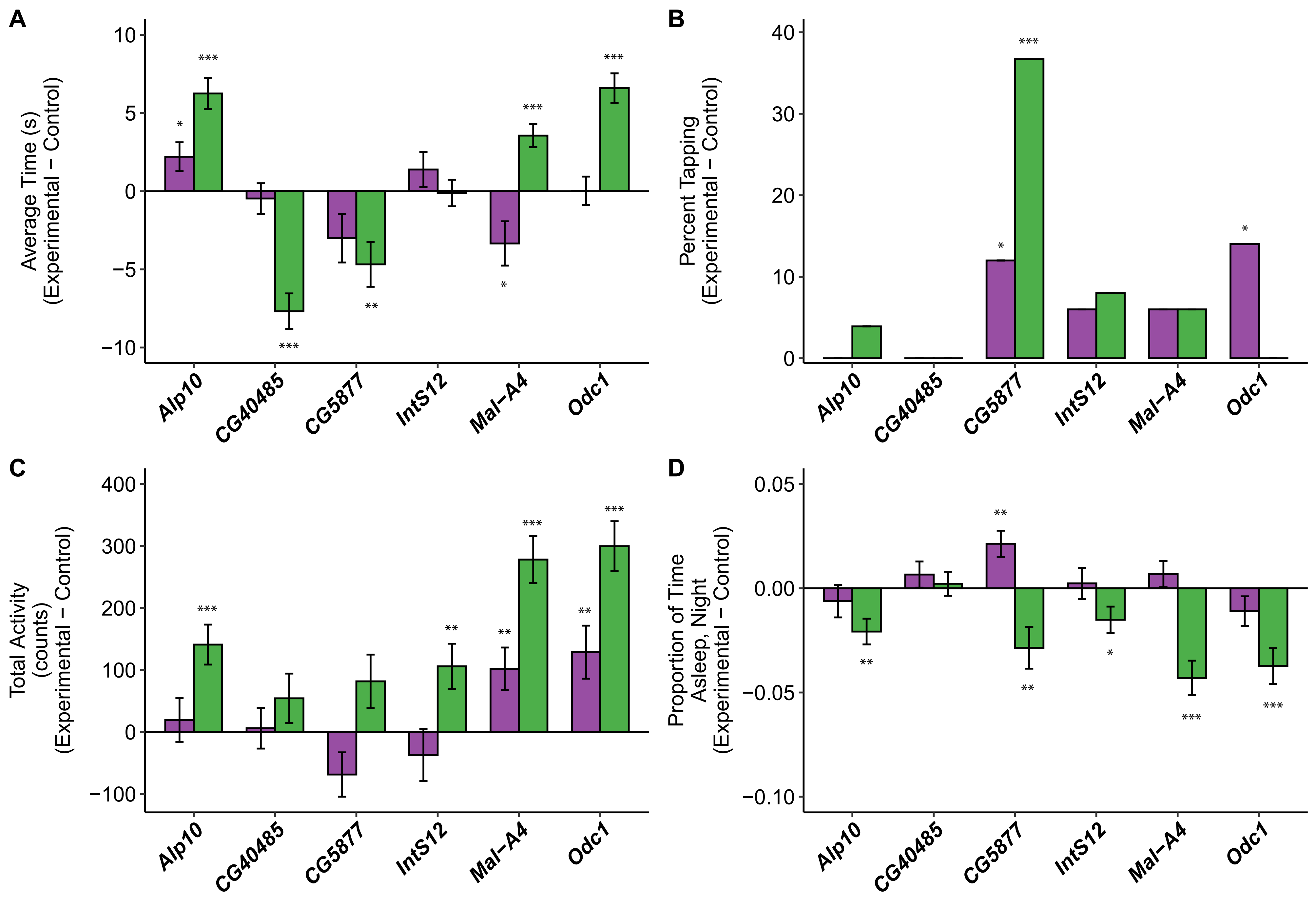
